# Time-restricted feeding exacerbates liver fibrosis by promoting BDH1-mediated ketolysis in hepatic stellate cells

**DOI:** 10.64898/2026.03.21.712927

**Authors:** Massimo Pinzani, Chang Pan, Wang Mingzhe, Patricia Lemnitzer

**Affiliations:** Department of Immunology, Ophthalmology and ENT, School of Medicine, Universidad Complutense de Madrid, Spain (J.M.G.-G.); Human Phenome Institute, Zhangjiang Fudan International Innovation Center, Fudan University, Shanghai, China; Department of Pharmacology, School of Pharmacy, Fudan University, Shanghai, China; Obstetrics & Gynecology Hospital of Fudan University, State Key Laboratory of Genetic and Development of Complex Phenotypes, and Institutes of Biomedical Sciences, Fudan University, Shanghai, China

**Keywords:** Time-restricted feeding (TRF), Liver fibrosis, β-hydroxybutyrate (BHB), BDH1, Hepatic stellate cells (HSCs), Ketolysis, Metabolic reprogramming

## Abstract

Time-restricted feeding (TRF) is widely considered metabolically beneficial, yet its impact on chronic liver disease progression remains poorly defined. This study investigates the effects of TRF on liver fibrogenesis. Using carbon tetrachloride (CCl_4_)-induced, bile duct ligation (BDL)-induced, and choline-deficient, L-amino acid-defined high-fat diet (CDAHFD)-induced murine models of liver fibrosis, we demonstrate that TRF consistently exacerbates fibrotic injury. Mechanistically, TRF induces the systemic elevation of the ketone body β-hydroxybutyrate (BHB). We identify the ketolytic enzyme 3-hydroxybutyrate dehydrogenase 1 (BDH1) as a critical mediator of this process within hepatic stellate cells (HSCs). BDH1 expression is markedly upregulated in activated HSCs, enabling these cells to metabolize BHB. This BDH1-dependent ketolysis redirects BHB-derived carbons into the tricarboxylic acid cycle, supplying acetyl-CoA and citrate to drive de novo lipogenesis and support a profibrogenic metabolic state. Both the genetic ablation of Bdh1 specifically in HSCs and the inhibition of hepatic ketogenesis successfully abolished the pro-fibrotic effects of TRF and exogenous BHB administration. Conversely, exogenous BHB alone was sufficient to recapitulate the exacerbated fibrotic phenotype observed with TRF. These findings reveal a context-dependent, detrimental role for TRF during chronic liver injury, driven by BDH1-mediated metabolic reprogramming in HSCs. Consequently, dietary interventions that elevate systemic ketone bodies should be approached with caution in the setting of active liver fibrosis.

## 1. Introduction

Liver fibrosis, characterized by the excessive accumulation of extracellular matrix (ECM), represents a common pathological endpoint of chronic liver injury from various etiologies, including toxins, metabolic disorders, and cholestasis^1^. If uncontrolled, fibrosis progresses to cirrhosis, liver failure, and hepatocellular carcinoma, posing a significant global health burden. The activation of hepatic stellate cells (HSCs) is a central event in fibrogenesis, driving the deposition of ECM components^2,3^.

Time-restricted feeding (TRF), a dietary intervention that confines food intake to a specific daily window without caloric restriction, has gained attention for its beneficial effects on metabolic health. It is known to improve insulin sensitivity, restore circadian rhythms, and reduce inflammation. Preclinical studies suggest TRF may alleviate metabolic dysfunction-associated steatohepatitis (MASH)^4^. Paradoxically, some evidence indicates that under specific pathological conditions, such as TRF might exacerbate liver damage. This highlights a complex, context-dependent role for TRF, where its impact may vary based on the underlying metabolic and disease state, particularly in the progression of liver fibrosis—an area not fully understood.

Ketone bodies, primarily β-hydroxybutyrate (BHB), serve as crucial alternative energy substrates during fasting or metabolic stress^5^. While their role in providing energy for extrahepatic tissues is well-established, their direct effects on non-parenchymal liver cells, especially HSCs, remain largely unexplored. The enzyme 3-hydroxybutyrate dehydrogenase 1 (BDH1), which catalyzes the interconversion of acetoacetate and BHB, is a key mediator of ketone body oxidation (ketolysis). Recent evidence points towards metabolic reprogramming, including altered lipid and ketone metabolism, as a critical driver of HSC activation and persistence. However, whether and how the elevated ketone bodies induced by TRF influence HSC biology and contribute to liver fibrosis is unknown.

In this study, we investigated the effects of TRF on the progression of liver fibrosis and elucidated the underlying mechanism. Contrary to the anticipated protective role, we found that TRF exacerbated fibrosis in multiple murine models (CCl_4_, BDL, and CDAHFD). We identified that TRF-induced hyperketonemia and the subsequent BDH1-mediated ketolysis in HSCs are pivotal in promoting a profibrogenic metabolic program. Genetic and pharmacological inhibition of this pathway attenuated fibrogenesis. Our findings reveal a novel, detrimental role of TRF in the context of chronic liver injury, mediated by HSC-specific ketone body metabolism, and challenge the notion of TRF as a universally beneficial intervention for liver disease.

## 2. Results

### 2.1 TRF promote HSC activation and liver fibrogenesis

To investigate the impact of time-restricted feeding (TRF) on liver fibrosis, we employed three well-established murine models: carbon tetrachloride (CCl_4_)-induced, bile duct ligation (BDL)-induced, and choline-deficient, L-amino acid-defined, high-fat diet (CDAHFD)-induced liver fibrosis. Mice in each model were subjected to either ad libitum feeding (Ctrl) or the TRF regimen.

Consistently across all three models, TRF treatment significantly exacerbated the progression of liver fibrosis. Histopathological analysis by Masson’s trichrome staining revealed a marked increase in collagen deposition in the livers of TRF-fed mice compared to Ctrl-fed mice (Fig. 1, left panel). Quantitative assessments corroborated these observations. The fibrotic area, measured as the Masson-positive area (%), was significantly greater in the TRF group (CCl_4_: p=0.0432; BDL: p=0.0073; CDAHFD: p=0.0478) (Fig. 1A-C). Biochemical measurement of hepatic hydroxyproline content, a direct indicator of total collagen, was also significantly elevated by TRF treatment (CCl_4_: p=0.0022; BDL: p=0.0011; CDAHFD: p=0.0114) (Fig. 1D-F). Furthermore, the mRNA expression level of α-smooth muscle actin (α-SMA), a key marker for activated hepatic stellate cells (HSCs), was significantly upregulated in the TRF groups (CCl_4_: p=0.0019; BDL: p=0.0010; CDAHFD: p=0.0011) (Fig. 1G-I). TIMP1 is a marker of liver fibrosis, mainly secreted by hepatic stellate cells. The TRF group also showed an elevated phenomenon. However, when we examined the indicators related to liver injury, we found that the changes in ALT and AST did not have a significant impact. In WT mice, TRF does not have an effect on the indicators related to liver fibrosis. Collectively, these data demonstrate that TRF promotes liver fibrogenesis in multiple experimental settings, as evidenced by enhanced collagen accumulation and HSC activation.

**Figure 1.**
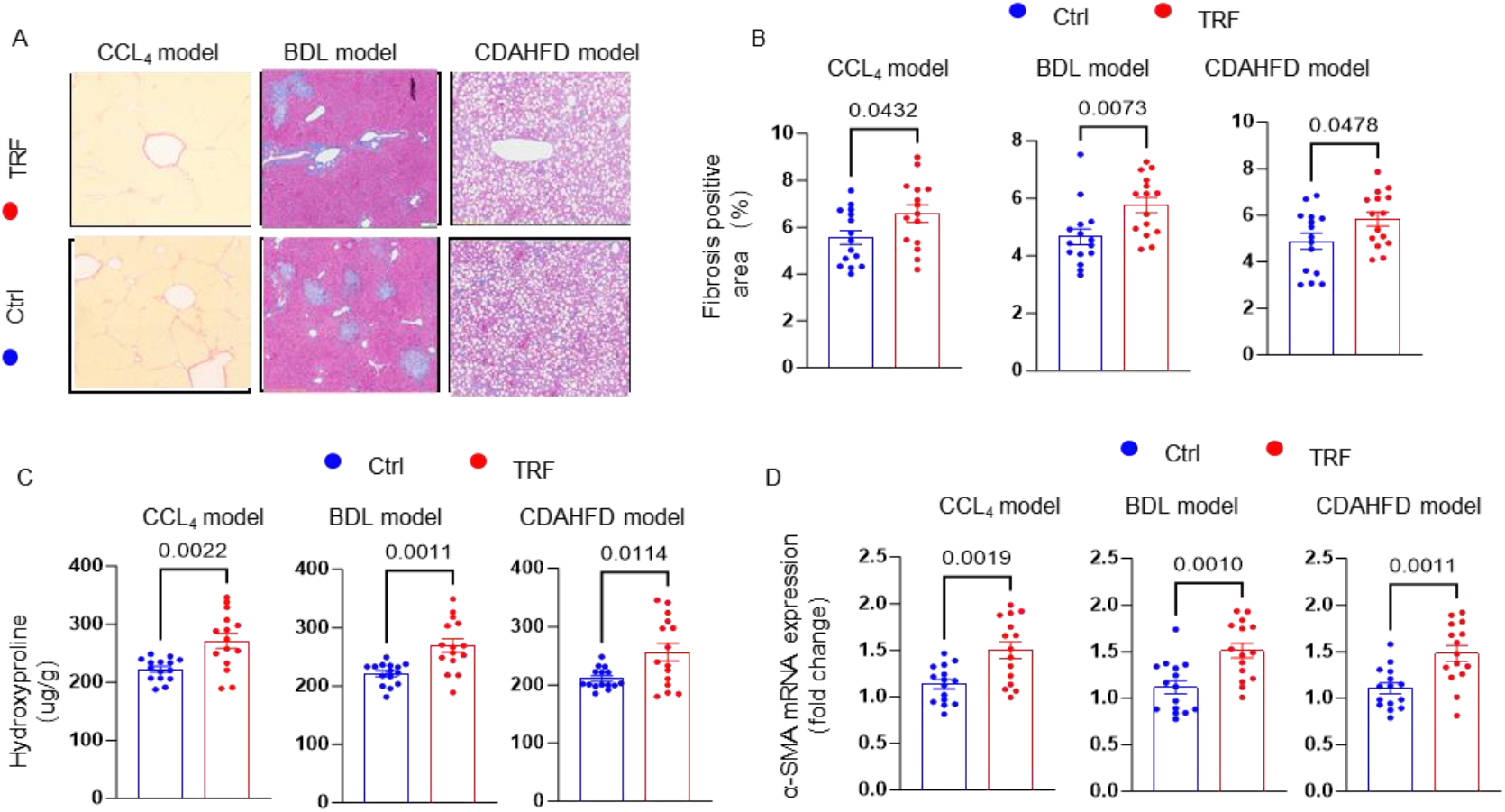
Time-restricted feeding (TRF) exacerbates liver fibrosis in multiple murine models. (A) Schematic representation of the experimental liver fibrosis models utilized: carbon tetrachloride (CCl_4_)-induced, bile duct ligation (BDL)-induced, and choline-deficient, L-amino acid-defined, high-fat diet (CDAHFD)-induced models. Mice were subjected to either ad libitum feeding (Ctrl) or TRF regimen. (B-D) Quantification of liver fibrosis in the three models. (B) Masson’s trichrome staining-positive area (%). (C) Hepatic hydroxyproline content, a biochemical marker of collagen deposition. (D) Relative mRNA expression levels of α-smooth muscle actin (α-SMA), a marker for activated hepatic stellate cells. Data are presented as mean ± SEM. Blue bars/scatter points represent the Ctrl group; red bars/scatter points represent the TRF group. Statistical significance was determined by unpaired two-tailed Student’s t-test (p< 0.05, *p< 0.01). Exact p-values are indicated on the graphs. TRF significantly promoted fibrosis progression across all models as evidenced by increased collagen deposition and activation of fibrogenic pathways.

### 2.2 TRF promotes HSC activation and liver fibrogenesis in a ketone body elevation-dependent manner in vivo

Given the consistent exacerbation of fibrosis by TRF, we investigated the underlying metabolic mediators. We hypothesized that enhanced ketogenesis might play a key role. Indeed, plasma levels of β-hydroxybutyrate (BHB), a primary ketone body, were significantly elevated across all three fibrotic models (CCl_4_, BDL, and CDAHFD) in TRF-fed mice compared to their ad libitum-fed counterparts (Fig. 2A).

**Figure 2.**
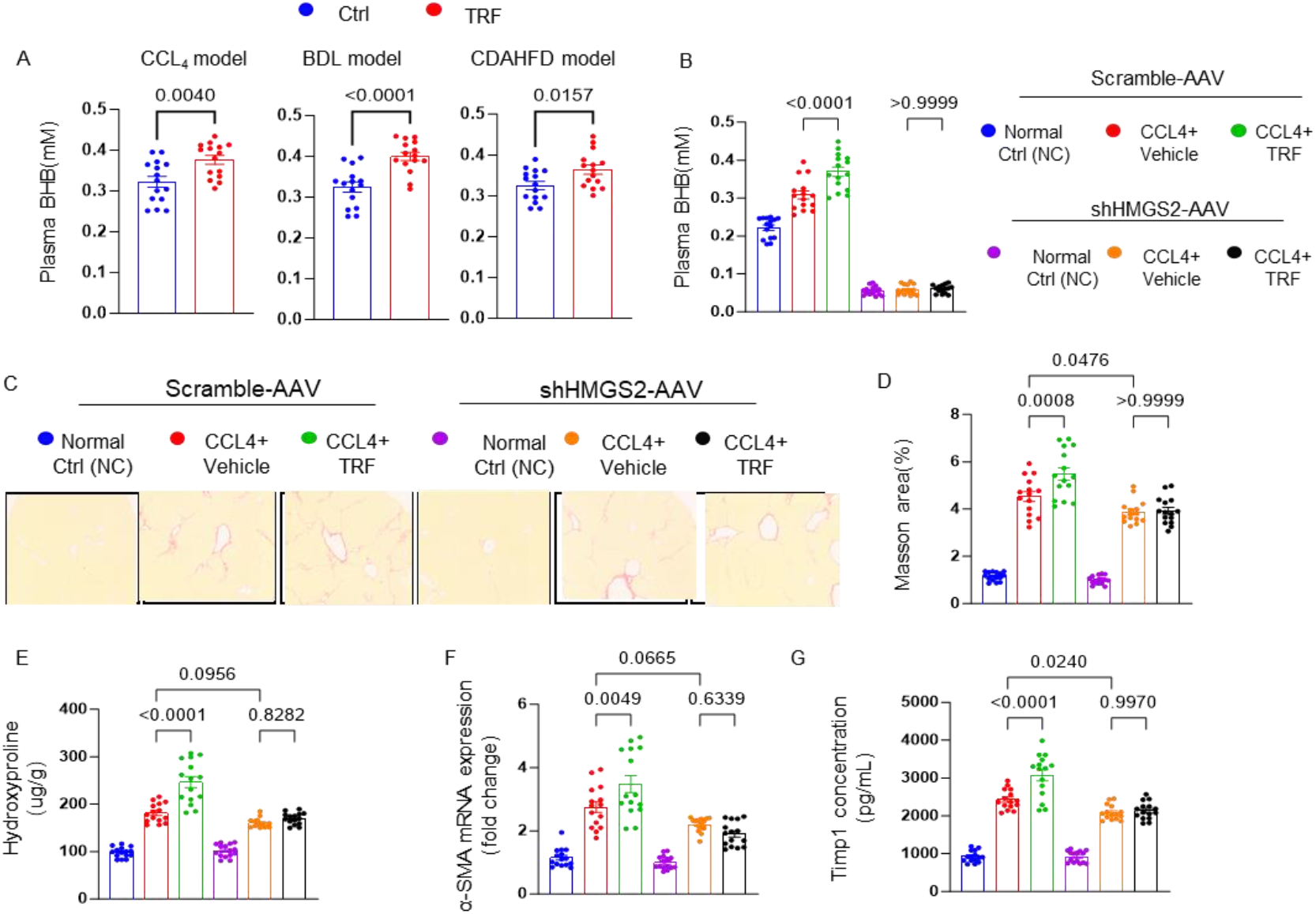
β-hydroxybutyrate (BHBA) is a key mediator of TRF-exacerbated liver fibrosis. (A) Plasma BHBA levels were elevated in TRF-fed mice subjected to CCl_4_, BDL, or CDAHFD models, compared to their respective ad libitumfed controls. (B) Plasma BHBA levels were significantly elevated in TRF-fed mice with CCl_4_-induced fibrosis, which was reversed by AAV-mediated hepatic knockdown of Hmgcs2(Hmgcs2-KD) but not by AAV-Control (AAV-Con). (C) [Description of panel C is intentionally omitted as the provided information indicates it is a blank panel]. (D-F) AAV-mediated hepatic Hmgcs2knockdown (Hmgcs2-KD) mitigated the TRF-aggravated fibrotic phenotypes in the CCl_4_ model. (D) Quantification of liver fibrosis area by Masson’s trichrome staining (%). (E) Measurement of hepatic hydroxyproline content. (F) Relative mRNA expression of α-SMA. (G) Plasma concentration of Timp-1 (Tissue inhibitor of metalloproteinase-1). Data are presented as mean ± SEM with individual data points overlaid on bar graphs. Blue bars/scatter points represent Ctrl group; red bars/scatter points represent TRF group. p< 0.05, *p< 0.01 (unpaired two-tailed Student’s t-test). AAV, adeno-associated virus.

To establish a causal link, we performed hepatic-specific knockdown of Hmgcs2, the rate-limiting enzyme for ketogenesis, using an adeno-associated virus (AAV) vector (shHmgcs2-AAV) in the CCl_4_ model. A control AAV (Scramble-AAV) was used for comparison. Hepatic Hmgcs2knockdown effectively abolished the TRF-induced increase in plasma BHBA (Fig. 2B). Strikingly, reversing the hyperketonemia also mitigated the pro-fibrotic effects of TRF. Hmgcs2 knockdown significantly attenuated the TRF-aggravated increase in collagen deposition, as quantified by both Masson’s trichrome staining area (Fig. 2D) and hepatic hydroxyproline content (Fig. 2E).

Furthermore, the upregulation of the activated hepatic stellate cell marker α-SMA mRNA in the TRF group was also blunted by Hmgcs2knockdown (Fig. 2F). Plasma levels of TIMP-1, a key regulator of extracellular matrix degradation, followed a similar trend, with TRF-induced elevation being reversed upon Hmgcs2 knockdown (Fig. 2G).

Collectively, these results demonstrate that TRF promotes a hyperketonemia state during liver injury and that genetic inhibition of hepatic ketogenesis is sufficient to abrogate the exacerbation of fibrosis, identifying BHB as a critical metabolic mediator of TRF-aggravated liver fibrogenesis.

### 2.3. BHB Promotes the Activation of HSCs In Vitro and in vivo

Having established ketone body elevation as a key mediator of TRF-aggravated fibrosis, we next sought to determine if β-hydroxybutyrate (BHB) acts directly on hepatic stellate cells (HSCs), the primary fibrogenic cell type in the liver. In vitro, treating primary mouse HSCs with exogenous BHB led to a significant and time-dependent increase in pro-collagen I production compared to normal control conditions. Furthermore, in the human HSC line LX2, BHB co-treatment potentiated the profibrogenic effect of TGF-β, a major fibrotic cytokine, as evidenced by a synergistic increase in α -SMA mRNA expression (Fig. 3B). These findings demonstrate that BHB can directly stimulate fibrogenic responses in HSCs.

**Figure 3.**
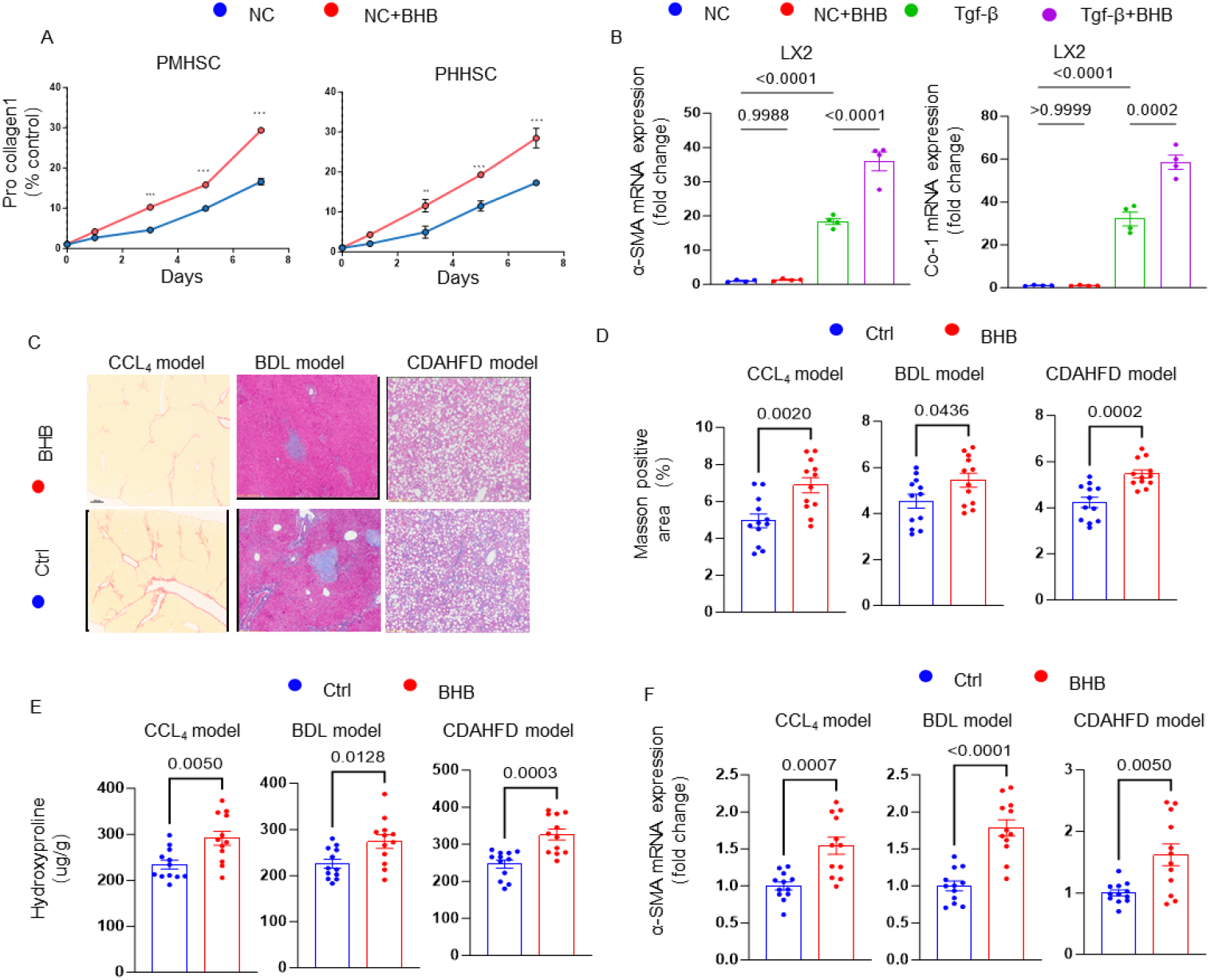
β-hydroxybutyrate (BHB) directly activates hepatic stellate cells and recapitulates TRF-exacerbated liver fibrosis in vivo. (A) Exogenous BHB treatment directly induces collagen synthesis in primary mouse hepatic stellate cells (PMHSCs). Cells were cultured under normal control (NC) conditions with or without BHB supplementation, and Pro-collagen I levels (% of control) were measured over time. (B) BHB amplifies the profibrogenic effect of TGF-β in the human hepatic stellate cell line LX2. α-SMA mRNA expression was assessed in cells treated with NC, NC+BHB, TGF-β, or TGF-β+BHB. (C-F) In vivoadministration of BHB mimics the pro-fibrotic effect of TRF in murine fibrosis models. Mice subjected to CCl_4_-, BDL-, or CDAHFD-induced fibrosis were treated with either vehicle (Ctrl) or BHB. (C) Schematic of BHB administration in the three models. (D) Quantification of liver fibrosis area by Masson’s trichrome staining (%). (E) Measurement of hepatic hydroxyproline content. (F) Relative mRNA expression of α-SMA. Data in A, B, D-F are presented as mean ± SEM. *p < 0.05, **p < 0.01 (unpaired two-tailed Student’s t-test for in vivo data; appropriate statistical test as performed for in vitro data).

To confirm the causal and sufficient role of BHB in vivo, we administered exogenous BHB to mice subjected to CCl_4_-, BDL-, or CDAHFD-induced fibrosis (Fig. 3C). Remarkably, BHB administration alone was sufficient to recapitulate the pro-fibrotic phenotype observed with TRF. BHB-treated mice exhibited a significant increase in liver fibrosis area by Masson’s trichrome staining across all models (Fig. 3D). Consistently, hepatic hydroxyproline content was elevated in the BHB group (Fig. 3E). Gene expression analysis further confirmed HSC activation, as shown by upregulated α-SMA mRNA levels (Fig. 3F). Together, these data provide direct evidence that BHB is not only necessary (as shown by Hmgcs2 knockdown) but also sufficient to drive liver fibrogenesis, acting as a primary effector molecule downstream of TRF.

### 2.4 BDH1-Mediated Ketone Body Metabolism Fuels Lipogenesis in Activated Hepatic Stellate Cells

To delineate the metabolic reprogramming driven by BDH1 in hepatic stellate cells (HSCs), we first mapped the potential pathway through which BDH1-mediated ketone body oxidation could fuel anabolic processes (Fig. 4A). We hypothesized that oxidation of β-hydroxybutyrate (BHB) generates acetyl-CoA, which can be utilized for de novo lipogenesis (DNL) after being exported from mitochondria as citrate.

**Figure 4.**
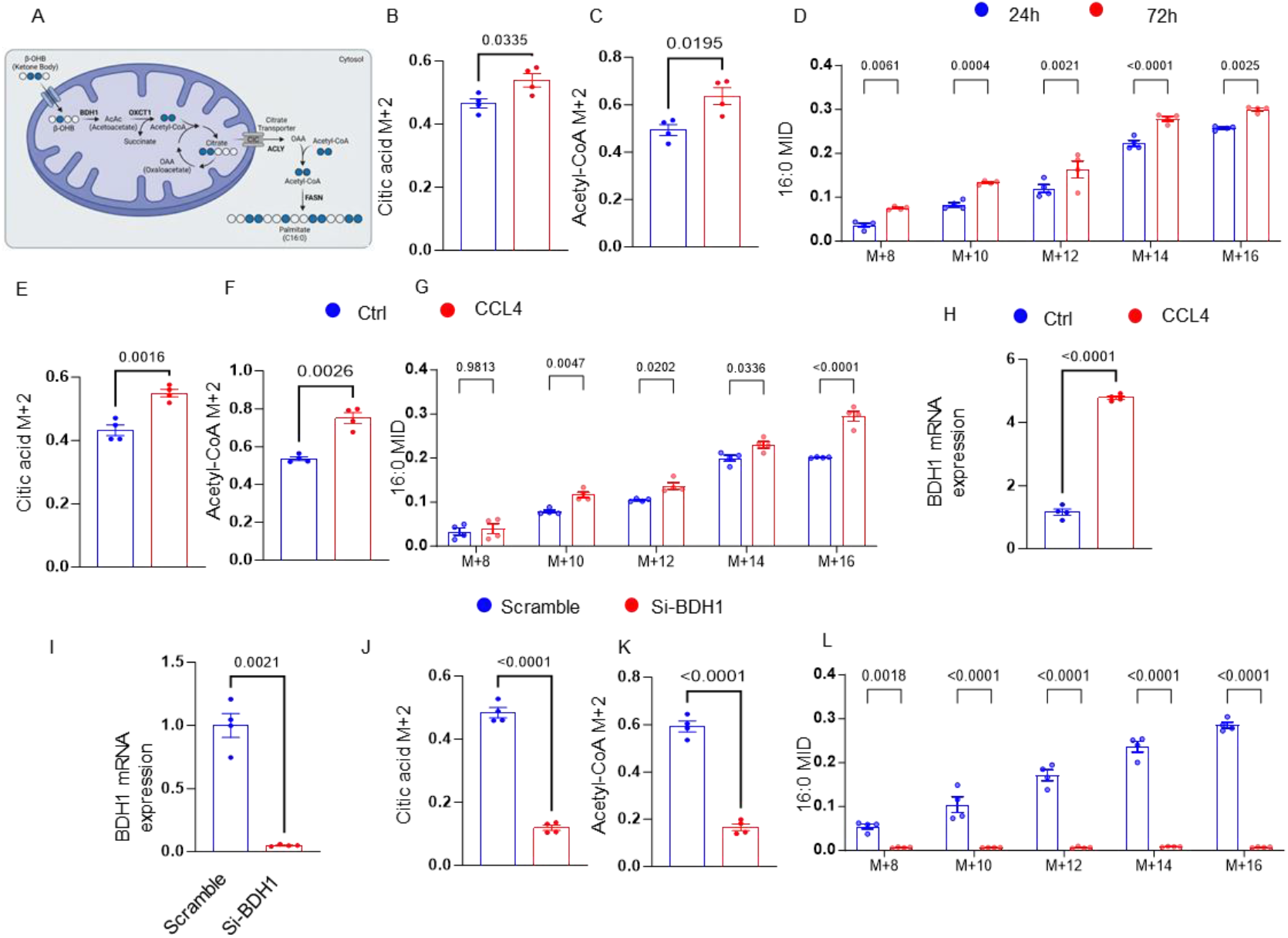
BDH1-mediated ketone body metabolism in hepatic stellate cells (HSCs) drives a profibrotic metabolic program. **(A)** Schematic depicting the central role of BDH1 in ketone body metabolism within HSCs. BDH1 catalyzes the interconversion of acetoacetate (AcAc) and β-hydroxybutyrate (BHB). Acetyl-CoA derived from BHB oxidation can fuel de novo lipogenesis (DNL) or enter the TCA cycle. **(B-D)** Stable isotope tracing reveals dynamic metabolic flux in primary human HSCs. HSCs were cultured with [U-13C]glucose, and metabolite enrichment was analyzed at 24h (blue) and 72h (red). **(B)** Molar percent enrichment (MPE) of citrate. **(C)** MPE of acetyl-CoA. **(D)** MPE of palmitate (16:0). Data indicate increased utilization of glucose-derived carbon for citrate synthesis and lipogenesis over time. **(E-F)** Comparison of key metabolites between control and fibrotic HSCs. **(E)** Relative levels of citrate. **(F)** Relative levels of AcAc and BHB. Fibrotic HSCs show altered ketone body and TCA cycle intermediate levels. **(G-H)** Western blot analysis confirms elevated expression of fibrotic markers and BDH1 in activated HSCs. **(G)** Protein levels of α-SMA and Collagen I. **(H)** Protein level of BDH1. **(1)**Validation of efficient BDH1 knockdown (BDH1-KD) in HSCs via siRNA, as shown by Western blot. **(J-M)** Functional consequences of BDH1 knockdown in HSCs. **(J)** Relative *BDH1*mRNA expression post-knockdown. **(K)** Intracellular BHB levels. **(L)** Intracellular citrate levels. **(M)** Expression of the fibrotic marker α-SMA (mRNA). BDH1-KD significantly reduces BHB and citrate levels and attenuates the expression of the fibrotic marker α-SMA. Data in bar graphs are presented as mean ± SEM. *p < 0.05, **p < 0.01**,*p < 0.001 (unpaired two-tailed Student’s t-test or one/two-way ANOVA as appropriate). AcAc, acetoacetate; BHB, β-hydroxybutyrate; BDH1, 3-hydroxybutyrate dehydrogenase 1; DNL, de novo lipogenesis; TCA, tricarboxylic acid cycle.

Using [U-^13^C]BHB tracing in primary human HSCs, we observed a time-dependent increase in the incorporation of glucose-derived carbon into key metabolic intermediates. The molar percent enrichment (MPE) of citrate (Fig. 4B) and its downstream product, acetyl-CoA (Fig. 4C), were significantly higher at 72 hours compared to 24 hours. This was concomitant with a pronounced increase in the mass isotopomer distribution (MID) of newly synthesized palmitate (16:0), indicating enhanced flux from glucose into fatty acid synthesis over time (Fig. 4D).

We next compared HSCs from fibrotic livers to control HSCs. Fibrotic HSCs exhibited a marked increase in citrate levels (Fig. 4E). Consistent with active ketone body metabolism, fibrotic HSCs also displayed elevated levels of citrate (Fig. 4J), acetyl-CoA and palmitate (16:0) (Fig. 4F), alongside increased protein expression of the fibrotic marker BDH1 itself (Fig. 4G, H).

To directly test the role of BDH1 in this metabolic shift, we performed BDH1 knockdown (BDH1-KD) in HSCs, which was confirmed at the protein level (Fig. 4I).Crucially, this was associated with a dramatic decrease in the levels of citrate (Fig. 4J) and acetyl-CoA (Fig. 4K). Furthermore, BDH1-KD resulted in a substantial suppression of de novo lipogenesis, as evidenced by a global decrease in the MID of newly synthesized palmitate (16:0) across all measured isotopomers (Fig. 4L).

Collectively, these data demonstrate that BDH1 is upregulated in fibrotic HSCs and is essential for maintaining elevated ketone body flux. This flux, in turn, supplies critical metabolites (citrate and acetyl-CoA) to fuel a lipogenic program, thereby identifying a novel metabolic axis that supports the activated, profibrogenic state of HSCs.

### 2.5 Genetic Ablation of BDH1 in Stellate Cells Attenuates TRF-Aggravated Liver Fibrosis

To determine if the pro-fibrogenic effects of BHB and TRF depend on BDH1-mediated metabolism within HSCs, we first examined the response of primary HSCs isolated from mice with a hepatocyte-specific deletion of Bdh1(Bdh1^f/f^; Lrat-Cre). In vitro treatment with BHB significantly increased the production of Pro-collagen I over time in HSCs from control (Bdh1^f/f^) mice. In stark contrast, HSCs lacking Bdh1 were completely resistant to this BHB-induced pro-fibrogenic stimulus, showing no increase in Pro-collagen I levels (Fig. 5A). A similar effect was observed for the activation marker α-SMA, whose mRNA expression was upregulated by BHB in control HSCs but not in Bdh1-deficient HSCs (Fig. 5B). These findings demonstrate that BDH1 expression in HSCs is essential for their fibrogenic response to BHB.

**Figure 5.**
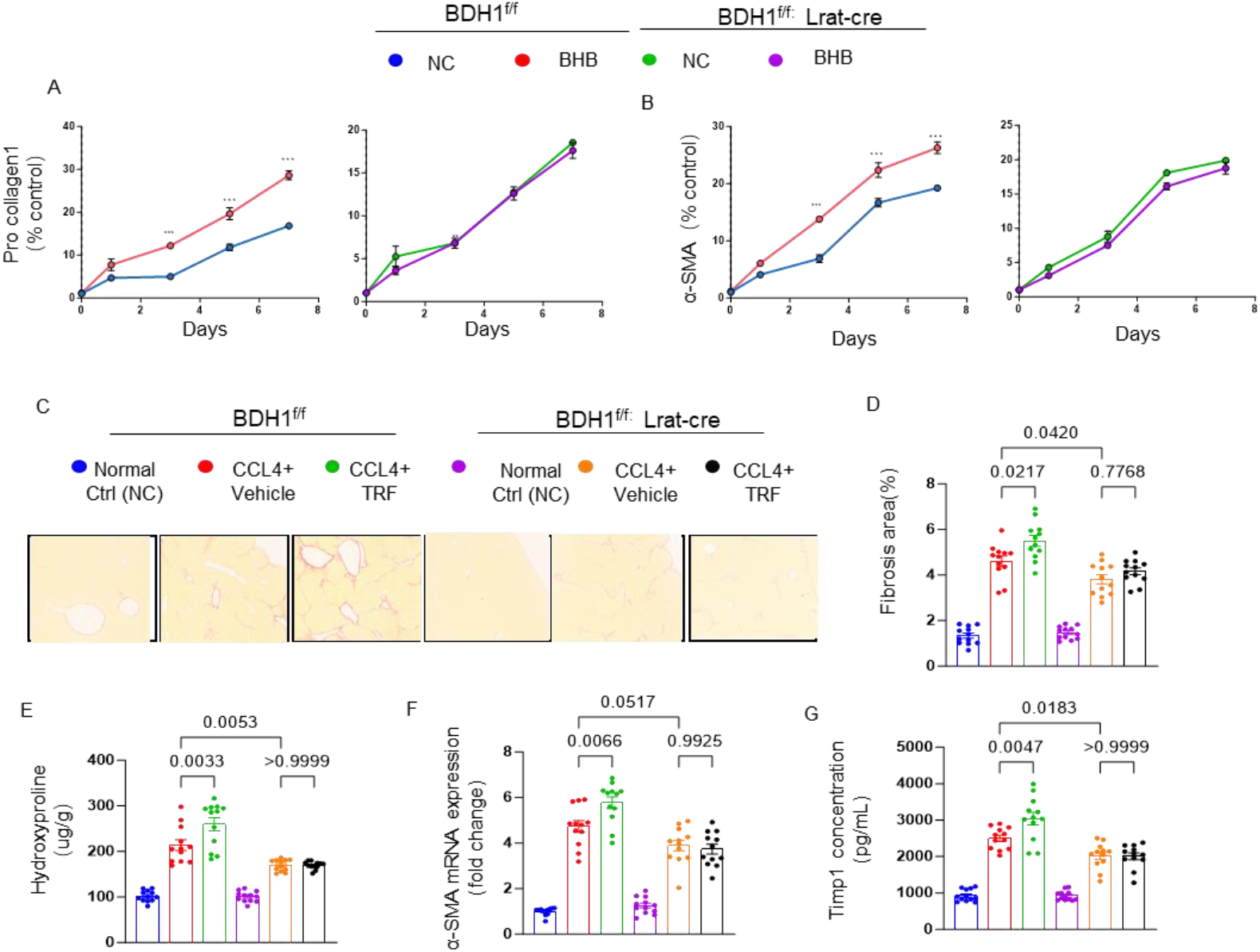
BDH1 in hepatic stellate cells is required for TRF-aggravated fibrosis. (A–B) A-B, Time-course analysis: Primary hepatic stellate cells (HSCs) isolated from BDH1fl/fl(control) and BDH1fl/fl;Lrat-Cre(HSC-specific BDH1knockout) mice were treated with or without BHB. Expression of the fibrosis markers (A) Pro-collagen1 and (B) α-SMA was measured over 8 days. Data are presented as percentage of control (% of BDH1fl/fl+ NC group at day 0). NC, normal control. (C-G), In vivo effects in a fibrosis model: BDH1fl/fland BDH1fl/fl;Lrat-Cremice were subjected to CCl4-induced liver fibrosis and treated with either vehicle or TRF. (C) Representative images of liver sections stained with Picro-Sirius Red (scale bar, 100 μ m). Quantitative analysis of (D) fibrosis area (%), (E) hepatic hydroxyproline content (a direct measure of collagen), (F) hepatic Acta2(α-SMA) mRNA expression by qRT-PCR, and (G) serum TIMP-1 concentration. NC, normal control group (no CCl4).

We next validated these observations in vivo using the CCl4-induced liver fibrosis model (Fig. 5C). As expected, CCl_4_ administration significantly increased all measured fibrosis parameters — including collagen deposition (Fig. 5D), hydroxyproline content (Fig. 5E), α-SMA expression (Fig. 5F), and plasma TIMP-1 levels (Fig. 5G)—in control mice (Bdh1^f/f^) compared to the Normal Ctrl group. Both exogenous BHB administration and the TRF regimen further exacerbated fibrosis in these control animals. Remarkably, HSC-specific knockout of Bdh1(Bdh1^f/f^; Lrat-Cre) provided significant protection. While Bdh1 deletion markedly attenuated the aggravating effects of both BHB and TRF. In Bdh1-knockout mice, the increases in collagen area, hydroxyproline content, α -SMA expression, and TIMP-1 concentration induced by TRF were all significantly reduced compared to treated control mice (Fig. 5D-G).

Collectively, these data establish that BDH1 activity in HSCs is a critical node required for the progression of liver fibrosis exacerbated by elevated ketogenic dietary pattern (TRF). Genetic ablation of Bdh1 specifically in HSCs renders them insensitive to the pro-fibrogenic signal of BHB in vitro and mitigates the worsening of liver fibrosis driven by metabolic.

## 3. Discussion

Although time-restricted feeding (TRF) is frequently proposed as a non-pharmacological treatment for metabolic disorders, including metabolic dysfunction-associated steatohepatitis (MASH), our results indicate that it exerts detrimental effects during chronic liver injury. Across toxin-induced (CCl_4_), cholestatic (BDL), and diet-induced (CDAHFD) murine models, TRF consistently worsened liver fibrogenesis. These data suggest that the physiological impact of TRF is disease-stage dependent, and its application may accelerate pathology in the context of active fibrosis.

The exacerbation of fibrosis associated with TRF correlates with systemic metabolic alterations, specifically the induction of hyperketonemia. While β-hydroxybutyrate (BHB) serves as an alternative energy substrate for extrahepatic tissues during fasting, our study identifies a profibrogenic function for BHB within the liver microenvironment. Exogenous BHB administration was sufficient to induce the fibrogenic phenotype observed in TRF-fed mice. Furthermore, inhibiting hepatic ketogenesis through Hmgcs2 knockdown prevented the TRF-induced exacerbation of fibrosis, confirming that elevated ketone bodies directly mediate this pathological response.

At the cellular level, the ketolytic enzyme 3-hydroxybutyrate dehydrogenase 1 (BDH1) drives this BHB-induced fibrogenesis in hepatic stellate cells (HSCs). Activated HSCs upregulate BDH1 to metabolize systemic BHB. This BDH1-dependent ketolysis redirects BHB-derived carbons into the tricarboxylic acid (TCA) cycle, generating citrate and acetyl-CoA to support de novo lipogenesis. This metabolic shift supplies the energy and intermediates required for HSC activation and collagen deposition. Hepatocyte-specific deletion of Bdh1 (Bdh1^f/f^; Lrat-Cre) protected mice from BHB- and TRF-induced fibrotic exacerbation, demonstrating that BDH1 activity in HSCs is strictly required for this process.

Our study has specific limitations. First, while the metabolic shift was consistent across three distinct murine models, the kinetics of TRF-induced ketosis and its fibrotic impact require validation in human cohorts. Second, the potential interactions between BHB-stimulated HSCs and other hepatic non-parenchymal cells were not fully explored and warrant subsequent investigation.

In summary, systemic ketosis induced by TRF promotes liver fibrogenesis through BDH1-mediated metabolic reprogramming in HSCs. These results indicate that dietary interventions requiring prolonged fasting or resulting in elevated ketone bodies should be evaluated with caution in patients with advanced liver disease. Targeting the BDH1 pathway in HSCs may offer a therapeutic strategy to restrict the metabolic substrates required for fibrotic progression.

## 4. Materials and Methods

### Animals and Liver Fibrosis Models

Mice were bred and maintained in groups of 5 animals per cage at the CNIC under specific pathogen-free conditions. Unless otherwise stated, males of 6-8 weeks old were used. Animal studies were approved by the local ethics committee. The local ethics committee approved all animal studies (PROEX 259/25; PROEX 345/46). Mice with HSC-specific deletion of Bdh1 (Bdh1^f/f^; Lrat-Cre) and wild-type littermates were also utilized. All animals were housed in a temperature-controlled room (22 ± 2°C) with a 12-hour light/dark cycle. CCl_4_ model: Mice were injected intraperitoneally (i.p.) with CCl_4_ (0.2 ml/kg) dissolved in corn oil, 3 times a week for 8 weeks. CDAHFD model: Mice were maintained on a choline-deficient, L-amino acid-defined, high-fat diet (Research Diets Inc.) for 8 weeks.

### In Vivo Interventions

To study the role of exogenous BHB, mice received intraperitoneal injections of sodium BHB twice daily at 12-hour intervals at a dose of 10 μmol/g body weight. Equimolar NaCl was used as the vehicle control. Gene Knockdown: Adeno-associated virus (AAV) vectors (shHmgcs2-AAV and Scramble-AAV) were utilized for hepatic-specific knockdown in vivo.

### Primary Cell Isolation and Culture

Primary HSCs: Mice were perfused via the portal vein with a 37°C EGTA solution, followed by in situ digestion using pronase (14 mg/35 mL, Millipore) and collagenase (20 mg/40 mL, Roche). Extracted livers were minced and further incubated in a shaking incubator with a pronase/collagenase solution containing 2 mg/mL DNase. The suspension was filtered through a 70 μm cell strainer, and HSCs were isolated via Nycodenze (Axies-Shield) density gradient centrifugation. HSCs were cultured in DMEM containing 10% FBS and Penicillin/Streptomycin. Primary HSCs were treated with 1 mM BHB starting on day 4 of culture and collected on day 7.

### Biochemical Analyses

Liver tissues were homogenized and hydrolyzed with 6 M HCl at 120°C for 22 hours. Hydrolysates were dried at 40°C for 72 hours, and absorbance was measured at 560 nm. Results were expressed as μg hydroxyproline per gram of liver tissue. Blood plasma was separated by centrifugation at 12,000 rpm at 4°C for 3 minutes. Plasma BHB levels were quantified using a BHB assay kit (Cayman Chemical).

### Histological Analyses

Livers were fixed in 4% paraformaldehyde for 24 hours and subjected to Sirius Red, or Masson’s trichrome staining.

### Western Blotting

Liver tissue samples were homogenized in RIPA lysis buffer (Thermo Fisher Scientific, Waltham, MA, USA) containing a protease inhibitor cocktail (Roche, Basel, Switzerland). Protein concentrations were determined using a BCA protein assay kit (Thermo Fisher Scientific). Equal amounts of protein were separated via 10% SDS-PAGE and subsequently transferred onto PVDF membranes (Millipore, Billerica, MA, USA). After blocking with 5% non-fat milk in TBST for 1 h at room temperature, the membranes were incubated with primary antibodies overnight at 4°C.

Protein bands were visualized using an ECL Western Blotting Detection Reagent (Cytiva, Marlborough, MA, USA).

### RNA Extraction and RT-qPCR

Total RNA was extracted from liver tissues using TRIzol reagent (Invitrogen, Carlsbad, CA, USA). The concentration and purity of the RNA were assessed using a NanoDrop 2000 spectrophotometer (Thermo Fisher Scientific). For cDNA synthesis, 1 μg of total RNA was reverse transcribed using the PrimeScript RT Master Mix (Takara Bio, Kusatsu, Japan). Quantitative real-time PCR (qPCR) was performed using TB Green Premix Ex Taq II (Takara Bio) on a QuantStudio 5 Real-Time PCR System (Applied Biosystems, Foster City, CA, USA). The relative expression levels of target genes were normalized to Gapdh as an internal control.

### Statistical Analysis

All data are expressed as the mean ± SEM. Statistical analyses were performed using GraphPad Prism software. Differences between two groups were analyzed using an unpaired Student’s t-test. For multiple groups, a one-way or two-way ANOVA was carried out, followed by a Bonferroni post hoc test. A value of p < 0.05 was considered statistically significant.

## Contribution

Patricia Lemnitze : Investigation, Writing-original draft, Data curation, Software, Methodology.

Chang Pan: Visualization, [Validation](https://zhida.zhihu.com/search?content_id=219282423&content_type=Article&match_order=1&q=Validation&zhida_source=entity)

Wang Mingzhe: Resources, Formal analysis

Massimo Pinzani: Conceptualization, Funding acquisition, Supervision, Writing – review & editing

